# Differentiating interactions of antimicrobials with Gram-negative and Gram-positive bacterial cell walls using molecular dynamics simulations

**DOI:** 10.1101/2022.07.15.500204

**Authors:** Rakesh Vaiwala, Pradyumn Sharma, K. Ganapathy Ayappa

**Affiliations:** Department of Chemical Engineering, Indian Institute of Science, Bangalore 560012, India; Centre for BioSystems Science and Engineering, Indian Institute of Science, Bangalore 560012, India; Eli Lilly Services India Private Limited, Bangalore 560103, India

## Abstract

Developing molecular models to capture the complex physicochemical architecture of the bacterial cell wall and to study the interaction with antibacterial molecules is an important aspect of assessing and developing novel antimicrobial molecules. We carried out molecular dynamics simulations using an atomistic model of peptidoglycan (PGN) to represent the architecture for Gram-positive *Staphylococcus aureus*. The model is developed to capture various structural features of the staphylococcal cell wall, such as the peptide orientation, area per disaccharide, glycan length distribution, crosslinking, and pore size. A comparison of the cell wall density and electrostatic potentials is made with a previously developed cell wall model of Gram-negative bacteria, *Escherichia coli*, and properties for both a single and multilayered structures of the Staphylococcal cell wall are studied. We investigated the interactions of the antimicrobial peptide melittin with the PGN structures. The depth of melittin binding to PGN is more pronounced in *E. coli* than *S. aureus*, and consequently the melittin has greater contacts with glycan units of *E. coli*. Contacts of melittin with the amino acids of peptidoglycan are comparable across both the strains, and the D-Ala residues, which are sites for transpeptidation, show enhanced interactions with melittin. A low energetic barrier is observed for translocation thymol with the four-layered peptidoglycan model. The molecular model developed for Gram-positive PGN allows us to compare and contrast the cell wall penetrating properties with Gram-negative strains and assess for the first time binding and translocation of antimicrobial molecules for Gram-positive cell walls.

## Introduction

Understanding the structure-function relations of bacterial cell envelopes enables development of potentially effective antimicrobial agents and therapies to curb virulent bacterial infections. Having structural diversity, the cell envelopes of Gram-positive bacteria are quite distinct from those of Gram-negative strains. The former possesses a 20-40 nm thick cell wall of peptidoglycans surrounding the cytoplasmic bilayer membrane,^1^ while the latter comprises of a more complex envelope with an asymmetric outer membrane of lipopolysaccharides and a thin periplasmic space protecting the cytoplasm. ^2^ Due to the structural diversity, the cell response and protective mechanisms are also distinct among the bacterial strains.

*Escherichia coli* (*E. coli*) and *Staphylococcus aureus* (*S. aureus*) are the typical strains that are representative of Gram-negative and Gram-positive bacteria, respectively. *S. aureus* is an opportunistic pathogen, causing nosocomial infections. With the emergence of multidrug resistant strains, such as Methicillin-resistant Staphylococcus aureus (MRSA), there is a pressing need to develop alternate treatment protocols. The cell wall plays a crucial role in infectivity and pathogenicity.^3^ Therefore, the Staphylococcal cell wall is a representative system of interest in clinical medicine for infections caused by Gram-positive strains.

Peptidoglycan (PGN) is one of the most essential constituents of cell wall of bacteria. It is an important target for antibiotics and antimicrobials. With its unique role as an exoskeleton, PGN resists turgor pressure within cells, dictates cell shape and is primarily responsible for structural rigidity of the cells.^4^ PGN is a mesh-like macromolecule and a heteropolymer consisting of amino-polysaccharides. The PGN precursors contain glycans, namely N-acetyl glucosamine (NAG) and N-acetylmuramic acid (NAM), found in uridine nucleotides, as well as several amino acids, hence justifies its name.^5,6^ Diamino acids such as diaminopimelic acid (m-A_2_pm) and lysine (Lys) are building blocks detected in the cell walls.^7–9^ Furthermore, the biochemistry of PGN confirms the presence of D-isomers of alanine (D-Ala) and glutamic acid (D-Glu), which are structurally distinct from regular amino acids.^10^

The architecture of PGN is the two- or three-dimensional polymer network comprising of glycan strands of NAG and NAM (Fig. 1). A disaccharide unit of NAG-NAM occupies ∼1 nm spacing along the axis of glycan helix,^11^ while the separation between the adjacent glycan strands in the network is ∼2 nm.^11,12^ A short stem of pentapeptide consisting of L- and D-isomers of alanine, glutamic acid and diamino acid covalently binds to the D-lactyl group of each muramic acid residue. The crosslinking among the glycan strands occurs through side chains in peptides, either directly via an amide linkage between amino group of diamino acid and carboxyl group of D-Ala in Gram-negative strains or through an interpeptide bridge in Gram-positive strains, as indicated in Fig. 1. We review some of the findings pertaining to the structure, architecture and dynamical interactions of cell wall of Gram-positive bacterium, specifically PGN of *S. aureus*.

**Figure 1:**
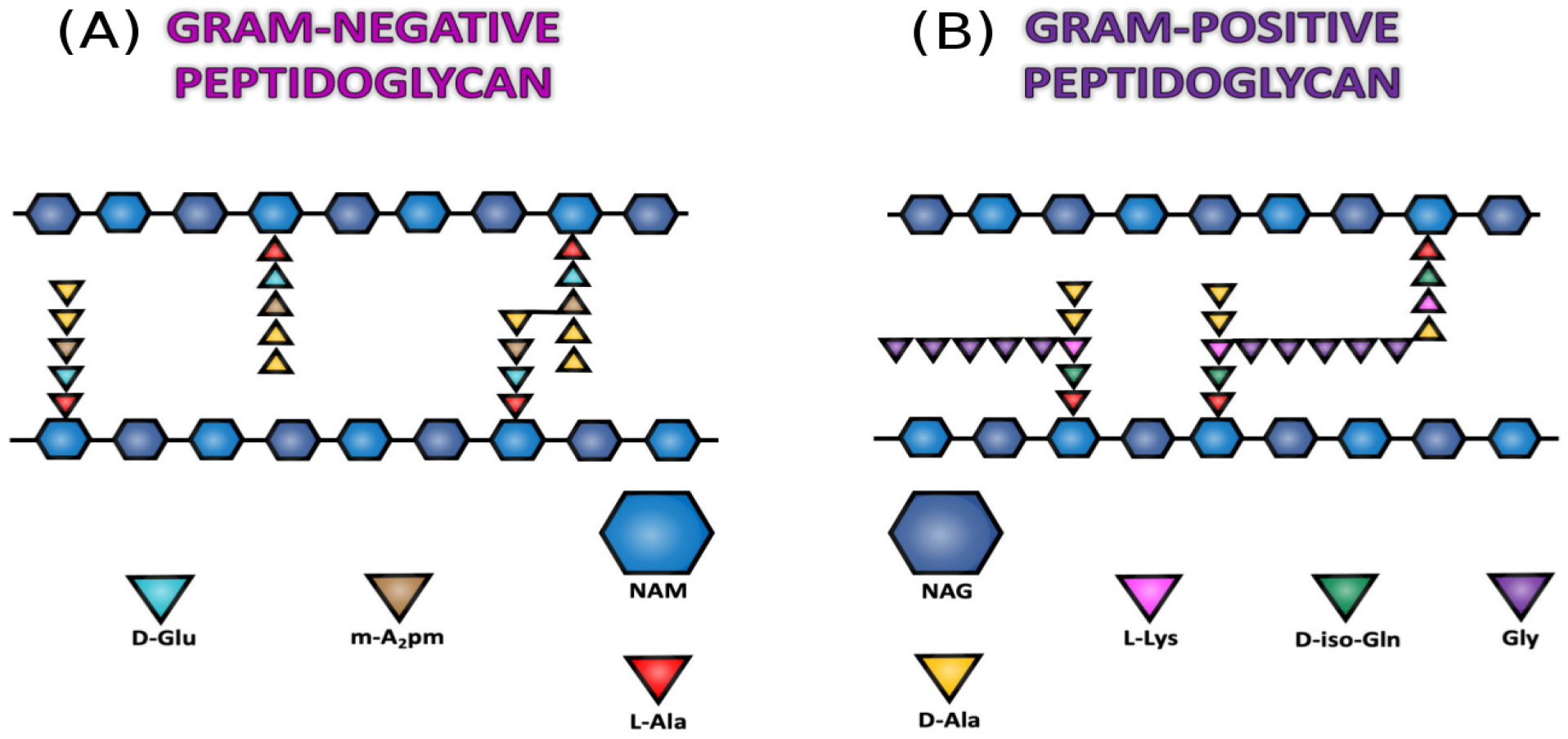
Schematics illustrating the molecular building blocks of peptidoglycans for (A) Gram-negative *E. coli* and (B) Gram-positive *S. aureus* bacteria. NAG and NAM sugars are shown by hexagons, while the amino acid residues in peptide stems are indicated by triangles. The crosslinking in *E. coli* occurs by direct bonding between m-A_2_pm and D-Ala residues, while the crosslinking in *S. aureus* involves a bridge of pentaglycine connecting L-Lys and D-Ala residues.

An experimental study^13^ using a reverse-phase high pressure liquid chromatography and mass spectrometry revealed that a major fraction of glycan strands of *S. aureus* comprises of only 3-10 disaccharide units of NAG-NAM, with an average length of 6 disaccharides. While 10-15% strands are longer than 25 disaccharides. X-ray diffraction study of isolated PGN and whole-cells *S. aureus* showed a four-fold helical orientational symmetry of the peptide stems.^11,14^ The peptide stems thus enable the crosslinking of glycans in two orthogonal directions. The interpeptide cross-link density was reported to be 67% using a solid-state NMR study.^15^ A study using cryo-electron microscopy confirmed an outer wall zone in cell envelope of *S. aureus*, consisting of a ∼19 nm thick and uniformly densed peptidoglycan-teichoic acid network.^16^ The isolated cell walls can expand and contract in response to salt and pH.^17^

Corroborating biosynthesis of interpeptide bridging in *S. aureus*, Kamiryo and Matsuhashi ^18^ identified that addition of a pentaglycine spacer to *ϵ*-amino group in Lys residues takes place sequentially from glycyl-tRNA, which acts as a glycine (Gly) donor. In a three-stage bridging process, the transferase FemX initiates incorporation of a Gly to Lys, followed by two more Gly using FemA. While the transferase FemB is essential for adding remaining two Gly to complete the bridge.^19^ The measured bridge-link density is 85%; this includes the cross-link density as well as the bridge-links that have C-terminal of pentaglycine bonded to Lys residues, while its N-terminal is not bonded to any other peptides^15^ (Fig. 1B). A tertiary cell wall structure of *S. aureus* was characterized in an experimental study of Sharif et al. ^20^ using CODEX spin diffusion of carbon-13. Accordingly, the peptide stems are in a plane perpendicular to the glycan chain, and the glycyl-carbonyl carbon of pentaglycine bridge lies within 5 Å from the anomeric carbon of the disaccharide. The three dimensional molecular models for PGN were designed in the work of Kelemen and Rogers.^21^

The contrasting views on orientation of cross-linked peptides were discussed in literature; these are parallel, anti-parallel and perpendicular orientations.^1^ The high degree of crosslinking, shorter glycan strands and longer bridges of pentaglycine in *S. aureus* support the parallel orientation viewpoint wherein the peptides that are cross-linked are parallel to each other. The crosslinking is restricted to only 50% in anti-parallel orientation. The cell walls with long glycan strands without interpeptide bridge (in *E. coli*) favour the cross-linked peptides orienting in the opposite directions. An architecture with cross-linked peptides orchestrated in perpendicular orientation is most suitable for the cell walls having intermediate bridge length and glycan chain length.^1^

A fragment of murein with disaccharide attached to decapeptide L-Ala–D-iso-Gln–L-Lys– (Gly)_5_–D-Ala–D-Ala was optimized in a computer simulation using molecular mechanics force field.^22^ In the proposed scaffold model, the glycan strands and oligopeptides take up the planar orientation, which is orthogonal to the cytoplasmic membrane. The peptides adopted helical conformations around the glycan strands. The simulated murein matrix resulted into 83% crosslinking, accommodating the experimental evidence of high degree of crosslinking in *S. aureus*.

To elucidate the architecture of cell wall of Gram-positive bacteria, electron cryo-tomography of murein of Bacillus subtilis showed a uniformly dense matrix with no internal structure.^23^ The study proposed a circumferential architecture of the PGN strands around the cell sacculus for rod-shaped Gram-positive bacteria, and provided an evidence using atomistic molecular dynamics simulations to support the proposed architecture.

The binding of antimicrobials with PGN inhibits transpeptidation and thereby impedes the PGN biosynthesis. In a simulation study on interactions of glycopeptides (vancomycin and its derivatives) with D-Ala–D-Ala termini of PGN precursors, the simplified models of *S. aureus* were developed.^24^ The energetic conformations of the glycopeptides were examined. Elucidating the structural insights into interactions of pentapeptide-pentaglycine stem of *S. aureus* with glycopeptide antibiotics, the studies using solid state NMR and molecular dynamics simulations revealed that the binding of Eremomycin to non-(D-Ala–D-Ala) segment of the peptide stem is facilitated by strong interactions of carboxyl terminus of the glycopeptide. While the N-methyl-leucine in the vancomycin is essential for strong binding to D-Ala–D-Ala termini. ^25,26^

To unravel the molecular mechanism of lysozyme mediated inhibition of *S. aureus*, a model lysozyme was docked with tetrasaccharide chain of NAG and NAM. ^27^ The latter was modified by O-acetylation at O6 in muramic residues. The structure of tetrasaccharide with O-acetylated modification was highly distorted, indicating high strain and instability in comparison with the chemically unmodified tetrasaccharide. Such structurally high energy conformations resulted in steric hindrances at the active cleavage site of lysozyme, leading to escape the sugar residues from lysing the glycosidic linkage. Perhaps the first detailed atomistic PGN model for *E. coli* was developed by Gumbart et al. ^28^ where a single layered PGN structure was studied. More recently we developed a MARTINI based model for the *E. coli* cell wall, and the free energy of interactions with thymol was assessed using umbrella sampling calculations.^29^ The interactions of surfactant molecules with a model PGN that mimics cell wall of *E. coli* has also been studied using molecular dynamics simulations. ^30^ The study reveals a correspondence between surfactant aggregation and their antimicrobial activity. Models for PGN with united atom descriptions have also been employed in simulation studies on the outer membranes of *E. coli*.^31,32^

It is clear from the above review that although the PGN precursors and simplified PGN models have been investigated in MD simulations for Gram-positive bacteria,^24–27^ a model that incorporates the naturally occurring three dimensional multilayered topology for the staphylococcal cell wall with its high degree of crosslinking is yet to be developed. In this work we develop an all-atom model of the staphylococcal PGN with the CHARMM36 compatible force field. The multilayered model constructed in this work incorporates the structural features of the *S. aureus* cell wall. In addition to the differences in cell wall thickness, differences in the amino acids in peptide stems as well as the manner in which peptides are cross-linked in Gram-positive and Gram-negative PGN models, differentiate between the Gram-positive and Gram-negative strains. We make a comparative assessment between the model cell wall structures of *S. aureus* and *E. coli*. We therefore also constructed single layered structures using PGN precursors of *S. aureus* and *E. coli*, and the structural properties were evaluated. Binding affinity of these structures with melittin peptides is assessed in the present study. The free energy for translocation of antimicrobial thymol molecules through the quadrilayered PGN structure is also computed using the densities of thymol molecules in a restraint free simulation.

## Molecular structures of peptidoglycans

In this section we describe the protocols used to construct the PGN strands and single layered structures for Gram-positive *S. aureus* as well as Gram-negative *E. coli* cell walls. We follow a similar methodology employed in previous studies^28,29^ to develop the single layered PGN model. In addition we also develop a more realistic multilayered PGN model for *S. aureus*. The molecular topology files for these model peptidoglycans are provided in the Electronic Supplementary Information (ESI).

### Single strands of peptidoglycans

Prior to building a complex three dimensional structure of PGN, we first constructed and simulated isolated PGN strands (Fig. 2A), which are basic precursors in a PGN network. An oligomeric PGN strand is comprised of a glycan backbone of NAG-NAM sugars and peptide stems of five amino acids attached to lactoyl carbon of muramic acid residues. The amino acids in PGN strands of *S. aureus* appear in a sequence L-Ala–D-iso-Gln–L-Lys–D-Ala– D-Ala, while those present in *E. coli* follow L-Ala–D-Glu–m-A_2_pm–D-Ala–D-Ala sequence (Fig. 1). Another difference between the PGN strands of *S. aureus* and *E. coli* is the presence of a reducing sugar at the terminal ends of the glycan strands in *S. aureus*, while the glycan chain in *E. coli* terminates with a non-reducing sugar. ^13^ Therefore, the glycan strands in *S. aureus* are modelled with a hydroxyl (‘OH’) group attached to the anomeric carbon of a NAM residue at the terminal ends. While, following our previous work,^29^ the strands in *E. coli* terminate with a methoxy (‘OCH_3_’) group bonded with anomeric carbon of NAM.

**Figure 2:**
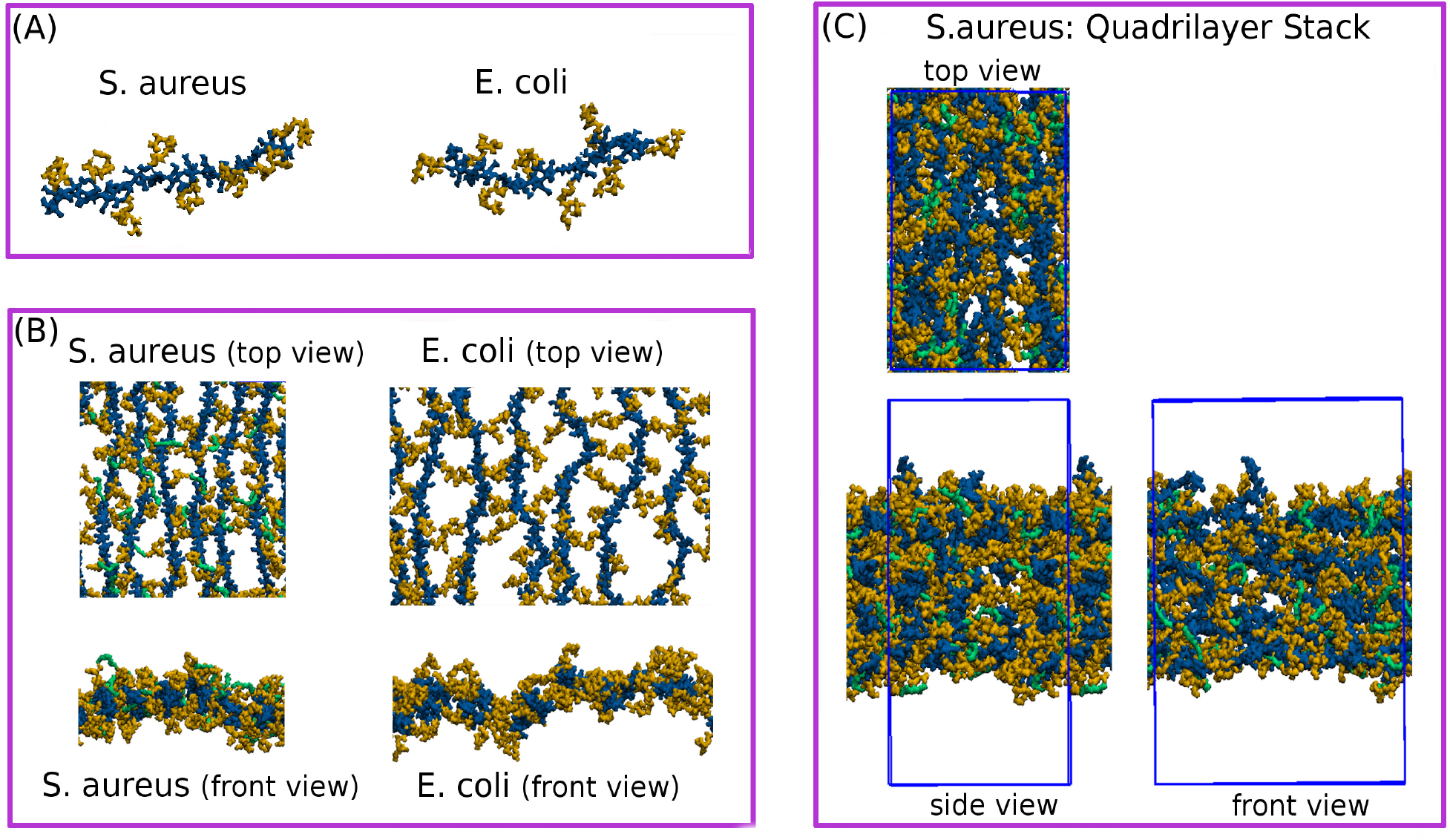
The model peptidoglycans for *S. aureus* and *E. coli* simulated in this study. (A) Single strands of peptidoglycan. The strands are 8-mer long each. (B) single layered models of peptidoglycans. (C) A four-layered stack of peptidoglycan for *S. aureus* cell wall. The color scheme refers to glycan strands (blue), peptide stems (orange), and pentaglycine bridges (green).

The model glycan strands are 8 disaccharide units long. The peptide stems in the model *S. aureus* are electrostatically neutral on account of the negatively charged D-Ala (−1e) and positively charged Lys residues (+1e). Each peptide stem in the *E. coli* model carries a net charge of -2e due to negative charges on Glu (−1e) and terminal D-Ala residue. These 8-mer strands (8 disaccharide unit long) of *S. aureus* and *E. coli* were simulated in a cubic box of size ∼11 nm. The system size and run time details are given in Table 1.

**Table 1:** System size and run-time details for the systems simulated in this study

| Systems | *S. aureus* |  | *E. coli* |  | Time<br>(ns) |
| --- | --- | --- | --- | --- | --- |
|  | PGN atoms | TIP3/Ions | PGN atoms | TIP3/Ions |  |
| Single PGN strand | 1051 | 42924 | 1054 | 52072/16(K<sup>+</sup>) | 400 |
| Single layered model | 12637 | 55333 | 11622 | 55375/182(K<sup>+</sup>) | 100 |
| Four-layered model | 26861 | 41580 | - | - | 200 |

### Single layered structures of peptidoglycans

The mesh-like networks mimicking the cell walls of *S. aureus* and *E. coli* (Fig. 2B) have been constructed using the PGN strands described above. The glycan strands are 13 disaccharide units long each, and periodically linked through glycosidic linkages at their terminal ends, to represent the infinitely long PGN chains. The peptides are cross-linked following the protocol reported in our earlier work.^29^ Initially a trial simulation was performed with the glycan strands weakly restrained using a harmonic force constant of 60 kJ mol^*−*1^ nm^*−*2^. The pairs of peptide stems which make contacts were identified with a distance cutoff of 0.85 nm over a trajectory of 50 ns. The peptides forming the contacts were preferentially cross-linked based on the contact frequencies. Since the PGN structures are single layered, we restricted the crosslinking to ∼50% for both the *S. aureus* and *E. coli* models. It is to be noted that the crosslinking in *E. coli* results in forming an amide bond between the nitrogen atom in the m-A_2_pm residue of an acceptor peptide and the carbonyl carbon of the D-Ala residue at position-4 in the conjugate peptide as illustrated in Fig. 1A. In the process of crosslinking, the terminal D-Ala drops out from the donor peptides. In contrast, the crosslinking in *S. aureus* is not direct; the amidation of peptides occurs via a short bridge of five units of Gly as depicted in Fig. 1B. While forming a bridge, a short stem of pentaglycine moiety covalently attaches to the nitrogen atom in the side chain of Lys, resulting into a bridge-link. A major fraction of the bridge-links, as high as 80%, completes the crosslinking by further amidation of the pentaglycine bridges at their free ends with the D-Ala residues of the neighboring peptide stems. In order to mimic an infinitely long two-dimensional sheet of PGN in the plane of the cell wall, the crosslinking is rendered periodic by placing covalent bonds between the peptide stems which fall near the periodic boundaries.

In the *S. aureus* model, we formed 29 bridge-links, of which 23 are cross-links amounting to ∼50% crosslinking. This maintains the ratio of cross-links to bridge-links close to ∼80% which is consistent with experimental data.^1,15,33,34^ The *E. coli* model has total 23 crosslinks formed by the direct bonding between m-A_2_pm and D-Ala residues among the contact making peptides. The equilibrium box dimensions are ∼12.5 × 12.5 × 11.3 nm for the *S. aureus* model, while the *E. coli* model is an asymmetric with the box size ∼17.7 × 11.5 × 8.6 nm on account of having distinct set of amino acids in the PGN network of *E. coli*.

### Multilayered staphylococcal peptidoglycans

The cell wall in Gram-positive bacteria is relatively thicker. For *S. aureus*, it is ∼19 − 33 nm thick, composed of multiple layers of peptidoglycans, taking a shape of three dimensional structure.^16,34^ We constructed a cell wall model for *S. aureus* using 32 glycan strands (196 disaccharide units) arranged in four layers as shown in Fig. 2C. Each layer consists of 8 glycan strands with a varying length of 4 − 8 disaccharide units. A trial simulation was carried out in order to preferentially create the dimeric-peptide links, as described before for single layered models. The peptide stems were cross-linked in three dimensions using pentaglycine bridges, generating 69 bridge-links (72%) and 58 cross-links (60%) consistent with experimental reports of the mesh-like topology of Staphylococcal cell wall. ^1,15,33,34^

### Simulation methodology

We carried out molecular dynamics simulations at 310 K temperature and 1 bar pressure using Gromacs^35^ (version 2018.2) with three dimensional periodicity. The positions of atoms were updated with the leap-frog algorithm with a time step of 2 fs. The simulations were performed using CHARMM36 force field with TIP3P water model.^28,36,37^ The force field parameters for thymol were obtained from the CHARMM General Force Field (CGenFF).^38^ The system temperature was controlled by a Nosé Hoover thermostat with a coupling time constant of 1 ps.^39^ A barostat of Parrinello Rahman with a compressibility 4.5 × 10^*−*5^ bar^*−*1^ was employed to maintain the pressure. ^40^ A coupling constant of 5 ps was set for the barostat. The single PGN strands were simulated in isotropic condition of pressure, while the networks of PGN, which comprised of several PGN strands, were first simulated using an anisotropic pressure coupling to fully relax the box dimensions during equilibration runs. In acquisition runs, the PGN networks were further simulated using semi-isotropic conditions of pressure, with compressibility 0 and 4.5 × 10^*−*5^ bar^*−*1^ in x-y plane (plane of glycans) and z direction, respectively. The hydrogen bonds were constrained using LINCS algorithm.^41^ A distance cutoff of 1.2 nm was used to compute van der Waals forces as well as the short ranged Coulomb interactions. The Lennard-Jones forces were made to smoothly vanish over the distance 1.0-1.2 nm by a force switch technique implemented in Gromacs. Potassium and Chlorine ions were used to neutralize the charged systems. The long range part of the electrostatic interactions was computed in reciprocal space using Particle-Mesh-Ewald.^42^

The trajectories were analysed in Visual Molecular Dynamics (VMD) software.^43^

## Structural Properties

In this section we discuss several structural properties of the model cell wall structures of Gram-positive *S. aureus*, and compare them with those of PGN model of Gram-negative *E. coli*. The structural properties of interest are the end-to-end distance for oligomeric PGN chains, orientational symmetry of peptides along the glycan chains, the equilibrium area per disaccharide, mass and charge density distributions, electrostatic potential, and the void distributions for the model networks.

### Chain end-to-end distance

The equilibrium chain length for a glycan strand is characterized by its end-to-end distance (*R*_e_). Figure 3A shows the distributions of *R*_e_ for the model glycan strands. Having identical repeating unit in their glycan backbones, the distributions for *S. aureus* and *E. coli* models show excellent overlap, although the PGN models have distinct amino acid residues. A mean value of *R*_e_ for the isolated and flexible PGN strands is calculated to be 5.2 ± 0.8 nm for 8 disaccharide unit long strands. The length of a disaccharide in cell wall networks is 1.03 nm, indicating that the glycan strands are quite stretched in the cell wall networks.^34,44,45^ The rigidity of glycan strands and peptide flexibility have a profound impact on the wall pores, which in turn have a bearing on the exclusion and transport of molecules through the cell walls.^17^

**Figure 3:**
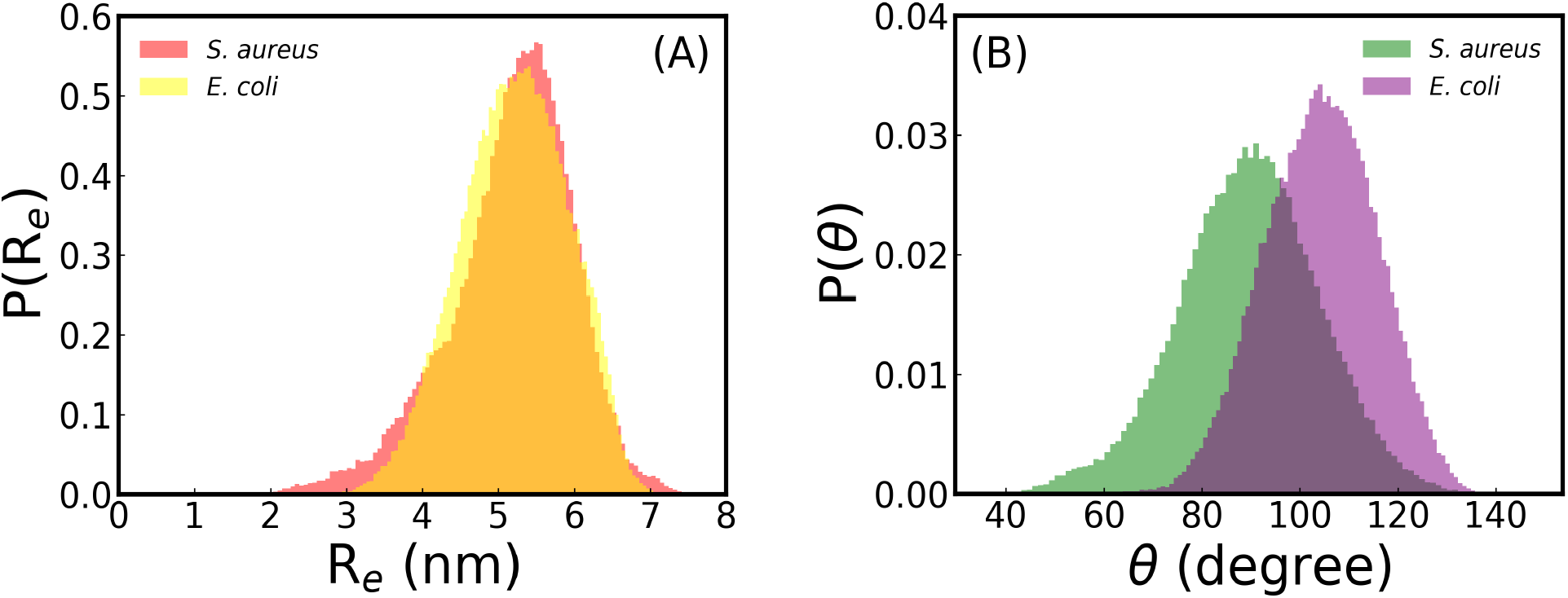
(A) Comparison of the end-to-end distances for the 8-mer PGN strands of *S. aureus* (−) and *E. coli* (−−) models. (B) Orientational angle between consecutive peptides along the PGN strands of *S. aureus* (−) and *E. coli* (−−). A vector connecting carbonyl carbon in lactoyl group of NAM and carbonyl carbon of terminal D-Ala residues is used to define the orientation of peptide stems, and θ is the angle between two consecutive orientational vectors of subsequent peptides along the glycan strands.

### Native periodicity of peptides

One of the factors that influences the extent of crosslinking in PGN networks is the orientation of peptides.^1^ In order to quantify the peptide orientation, we calculated vectors, each pointing from the lactoyl carbon of NAM residue to the terminal carbonyl carbon in D-Ala residue of a peptide stem.^28^ The equilibrium angle between these vectors across two consecutive peptides signifies a kind of symmetry of peptide orientation. The peptide stems along the glycan backbone adopt a four-fold symmetry of arrangement as evident in Fig. 3B. The mean orientational angles are 89.5° ± 14.5° and 104.2° ± 11.5° for the 8 disaccharide unit long glycan strands of *S. aureus* and *E. coli* models, respectively. A fact that four-fold periodicity of peptide orientation enables higher degree of crosslinking in three dimensional PGN networks is consistent with the studies on the *S. aureus* cell walls.^15,33^ In contrast, only half of the peptides in theory can form cross-links for the in-plane structures, resulting in a low percentage of crosslinking for the *E. coli* cell walls.^1,44,46,47^

### Surface area

For the single layered PGN models, the length of glycan backbones is calculated to be ∼0.95 nm/disaccharide unit, which is consistent with 1.03 nm reported earlier.^34,44,45,48^ From equilibrium lateral dimensions, the surface area per disaccharides turns out to be ∼1.72 nm^2^ and ∼2.24 nm^2^ for the *S. aureus* and *E. coli* models, respectively. The relatively higher value of area per disaccharide for the single layered *E. coli* is attributed to the electrostatic repulsions among the negatively charged peptide stems. Furthermore, the surface area for the *E. coli* model is consistent with ∼2.5 nm^2^ from electron microscopy data.^49^ This difference in area per disaccharide will have a direct impact on pore size distributions in the cell walls as well as the permeability of molecules.

### Density profiles

For the out-of-plane structure of the cell wall, the distribution of glycans and peptides in a direction normal to the plane of PGN provides an estimate of thickness of the PGN layer. Figures 4A and D illustrate the mass density of PGN across the z-direction for the single layered *S. aureus* and *E. coli* structures respectively, which are laterally extended in other orthogonal directions. In comparison to the *S. aureus* model, the mass density distribution for *E. coli* is broader with a thickness of ∼4.8 nm calculated at 10% of the peak density. The thickness of the *E. coli* model is consistent with previous simulation studies.^28,29^ Another distinguishing feature arising from the differences in the area per disaccharide of Grampositive *S. aureus* and Gram-negative *E. coli* is the water density at the core of cell wall. A relatively lower water density in the *S. aureus* structure indicates a lower affinity for water in the *S. aureus* cell wall when compared with *E. coli*, which has higher area per disaccharide and the wider pore size distribution (Figs. 4I and J). Potassium ions, which are used to neutralize the charges on PGN structure of *E. coli*, are symmetrically distributed across z-direction with the peak concentration at the core of the structure as indicated in an inset in Fig. 4D.

**Figure 4:**
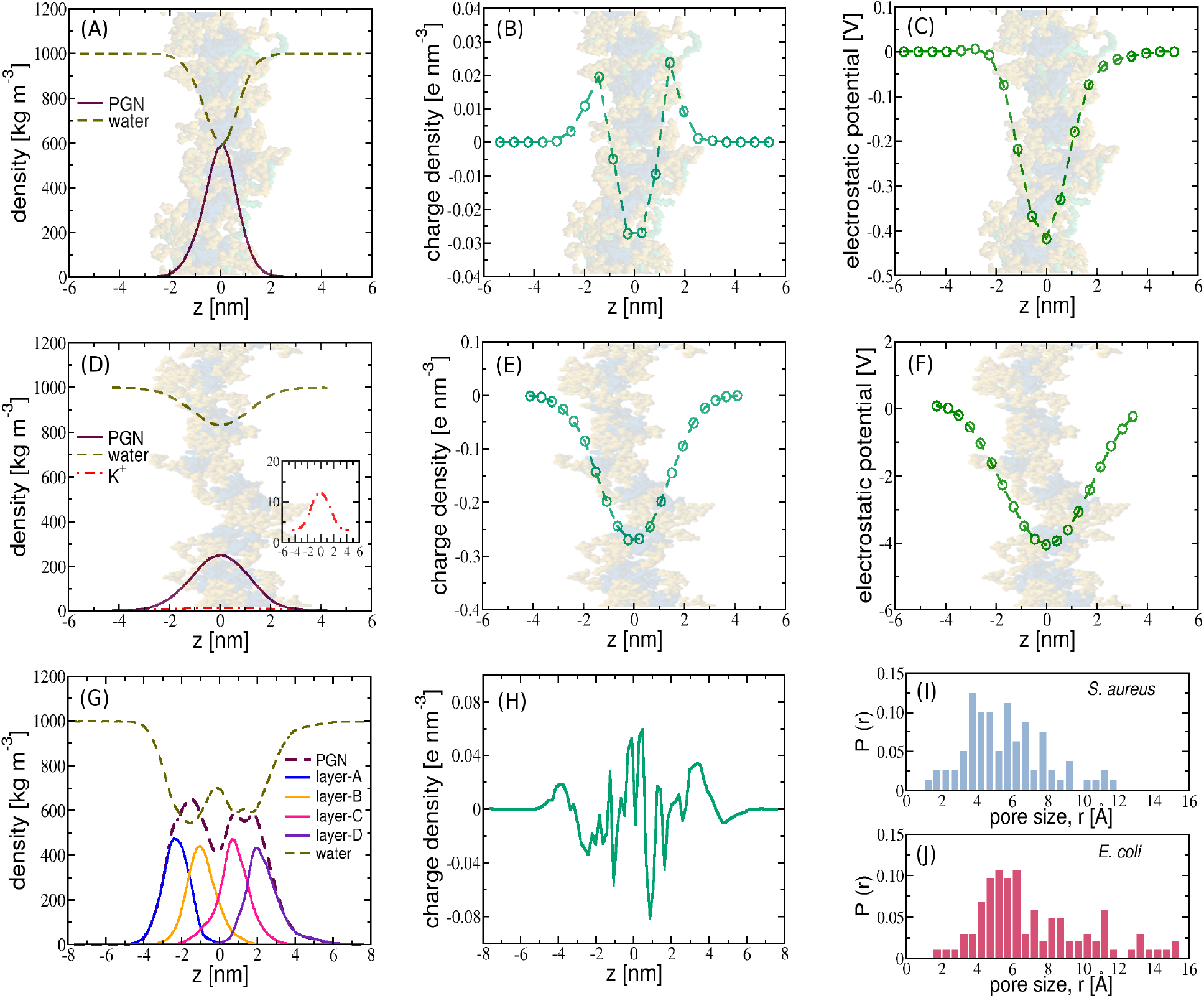
The structural profiles and electrostatic distributions for the single as well as multilayered models of cell walls. The top panel shows the distributions for (A) mass density, (B) charge density and (C) electrostatic potential across z-direction (normal to the plane of PGN) for the single layered *S. aureus* model. The corresponding distributions for *E. coli* model are shown in the central panel (D-F), while the panel at bottom indicates (G) mass density and (H) charge density for the four-layered model of *S. aureus*. The translucent images illustrate the peptidoglycan sheets. The histograms for void size distributions are compared for single layered models of (I) *S. aureus* and (J) *E. coli*.

For the quadrilayer stack of *S. aureus* PGN, the density of PGN shows four distinct peaks, corresponding to four PGN layers (Fig. 4G). The widths of the distributions are nearly similar for all four layers, indicating the uniformity in size of the multilayered topology of PGN. The porous structure has cavities with a low mass density of PGN in the regions ranging over −1 ≤ *z* ≤ 1 nm. The water density in these cavities is relatively high and it is manifested as a peak in water density at z=0.

### Voids in peptidoglycan structures

We calculated the pore size distributions for the single layered PGN structures of *S. aureus* and *E. coli* using the method described in our previous work.^29^ As depicted in Fig. 4I, the pore size distribution for the model *E. coli* structure is wider with pores as large as 1.5 nm. The pores with the higher densities are in the size range 0.4-0.8 nm. The maximum pore size for the model *E. coli* is consistent with the pore radius of ∼2 nm reported in the previous experimental as well as simulation works.^28,50^ The higher water content observed in the core of the PGN structure of *E. coli* (Fig. 4D) is a direct consequence of having larger size pores in the PGN of *E. coli*. The electrostatic potential is a plausible cause for driving water dipoles into the region of high potential.

### Electrostatics

Membrane disrupting property of antimicrobial peptides is solely governed by the electrostatic interactions.^51^ Having differences in the charged residues in amino acid precursors of PGN, it is important to bring out the contrasting features in electrostatics profiles for the model PGN structures of *E. coli* and *S. aureus*. Figures 4B and E show the distributions of charges along a direction orthogonal to the plane of PGN for the model single layered structures. The charge density is as low as -0.03 e nm^*−*2^ for *S. aureus*, while it is nearly ten-fold higher in magnitude for *E. coli* model. The reason for the comparatively higher negative charge density for the PGN of *E. coli* is the presence of anionic charges on *E. coli* peptides compared to the neutral peptides in PGN structure of *S. aureus*. This difference in the charge densities is also apparent in the electrostatic potentials given in Figs. 4C and F. The magnitude of potential is proportionately higher for the *E. coli* model compared to *S. aureus*. On account of high negative potential, the enhanced electrostatic repulsions among the peptides in the *E. coli* structure cause the interstrand spacing between the glycan strands wider, resulting into an increased area per disaccharide for the *E. coli* structure compared to the single layered structure of *S. aureus* as mentioned before. The charge density profile for the multilayered PGN model of *S. aureus* is rugged with crests and troughs (Fig. 4H).

### Interactions of peptidoglycans with melittin

The single layered models of PGN discussed in previous sections have been employed to investigate the interactions of PGN precursors with melittin peptides. Melittin is a naturally occurring antimicrobial peptide (PDB entry: 2MLT). Melittin is comprised of 26 residues, with polar and non-polar amino acids unevenly distributed. It is a prominent component in the venom of honey bees (*Apis mellifera*), and it has been widely investigated as a antimicrobial agent against bacteria and fungi.^52–54^ Melittin is recognized as a promising antibacterial agent in healing MRSA-infected skin wounds.^55,56^ Melittin tends to lose its native helical structure in water (Fig. S1 in ESI). This unfolding of melittin in aqueous environment has been reported previously in molecular dynamics simulation studies.^57^ Using transmission electron microscopy, a study on melittin treated *S. aureus* cells showed the morphological changes with cytoplasmic disintegration. ^58^ Our recent study reports the free energy landscapes for CM15 peptide (cecropin A and melittin residues) for its translocation across the complex outer membrane of *E. coli*.^59^

Although the interactions of melittin with lipid membranes have been extensively studied and several molecular insights have emerged^60–64^ only few studies have reported the interaction with the more complex components of the bacterial cell wall. Using single-molecule fluorescence microscopy, the electrostatic interactions of melittin with lipid A containing supported bilayers have been investigated, and melittin showed a much slower diffusion when bound to the lipid A moieties compared to that of the phospholipid-bound melittin. ^65^ The efficacy of melittin in disruption of biofilms formed by *S. aureus* and *E. coli* has been investigated using experimental assays, and it has been found that the peptide is more effective against biofilm formation by *S. aureus* than *E. coli*.^66^

A molecular understanding of cell wall interactions with antimicrobial peptides remains elusive.^67^ To bring out detail insights on binding of melittin peptide with model PGN structures, we have simulated the single layered PGN models with 8 melittin peptides (helix-turn-helix structure) initially placed in bulk water. We also carried out simulations with unfolded melittins interacting with PGN models to study the possibility of structural transitions, if any, within 500 ns of simulation runs. The system details are given in Table 2. The lateral dimensions of the model cell walls were ∼12.5 × 12.5 nm and ∼17.5 × 11.5 nm for the *S. aureus* and *E. coli* structures, respectively.

**Table 2:** Model cell walls interactions with small actives

| Systems | PGN atoms | TIP3/Ions | Actives |
| --- | --- | --- | --- |
| Single layered model (*S. aureus*)<sup>a</sup> | 12637 | 53851/40(Cl<sup>-</sup>) | 8 (Melittin) |
| Single layered model (*E. coli*)<sup>a</sup> | 11622 | 73487/142(K<sup>+</sup>) | 8 (Melittin) |
| Single layered model (*S. aureus*)<sup>b</sup> | 12637 | 52558/20(Cl<sup>-</sup>) | 4 (Melittin) |
| Single layered model (*E. coli*)<sup>b</sup> | 11622 | 71736/162 (K<sup>+</sup>) | 4 (Melittin) |
| Four-layered model (*S. aureus*) | 26861 | 39076 | 30 (Thymol) |

<sup>a</sup> melittin in native state
<sup>b</sup> melittin in unfolded state

### Melittin binds strongly with *E. coli*

Figures 5A and D show the time evolution of z-coordinates of the center-of-mass of melittin molecules interacting with model *S. aureus* and *E. coli* structures. Prone to interact favourably with PGN, melittin eventually approaches the PGN during the course of a 1 *µ*s simulation (Fig. 6). The depth of peptide penetration is stronger for the *E. coli* model when compared with interactions with the *S. aureus* PGN. This increased propensity toward the PGN of *E. coli* can be seen in Figs. 5B and E where a relatively higher density of melittins in the core of the *E. coli* structure is observed and all the melittin molecules are well within the PGN layer of *E. coli*. In the case of *S. aureus* the melittin molecules are more spread out across the membrane and we observe a greater number of surface bound peptides.

**Figure 5:**
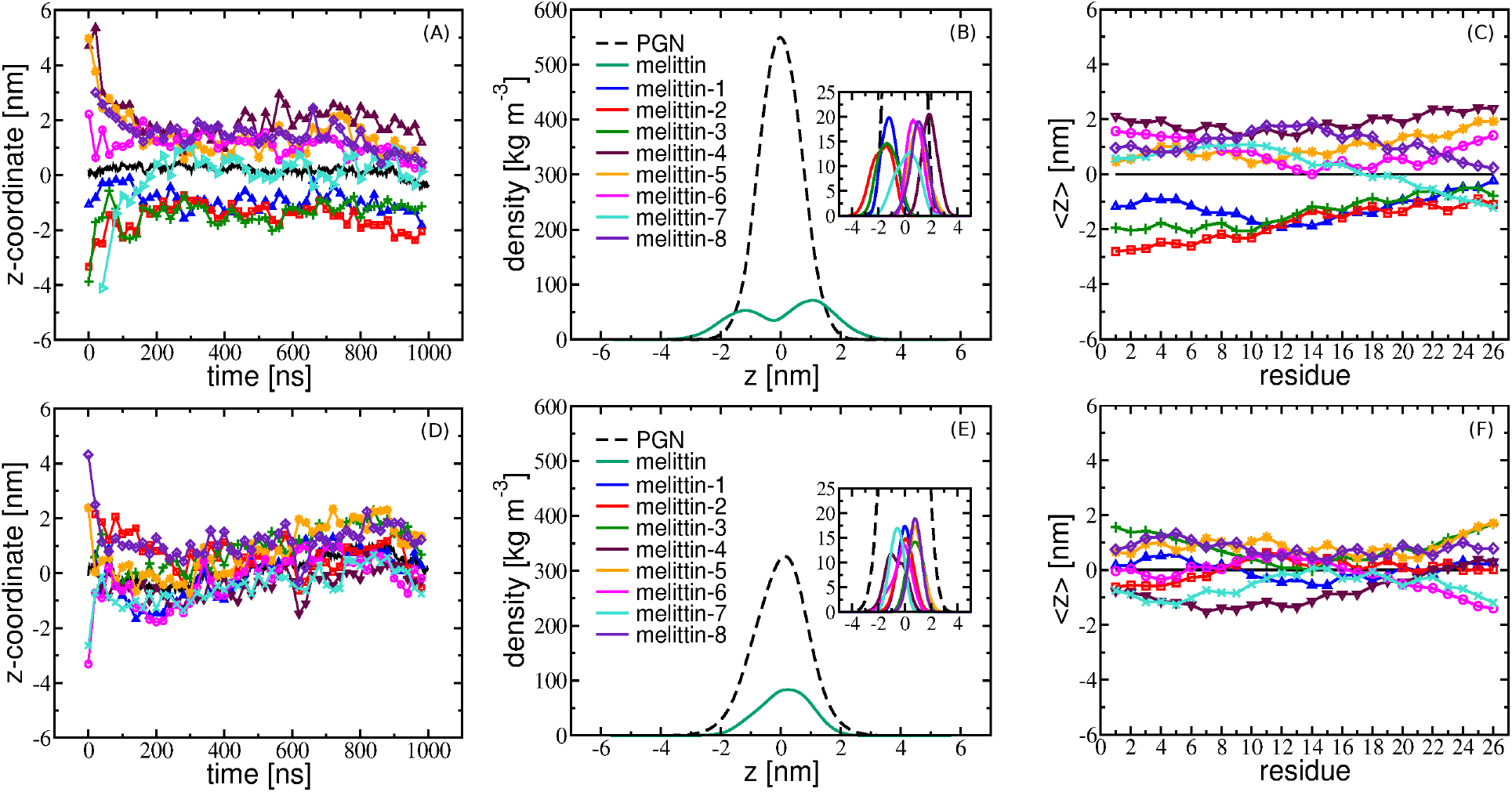
The results from melittin simulations indicating the depth of interactions of melittin with peptidoglycans. The top panel corresponds to the model *S. aureus*, while the bottom panel refers to the *E. coli* model. (A, D) The z-coordinate of the center-of-mass of peptidoglycans as well as individual melittin. (B, E) Density profiles for PGN and melittin peptides in the direction orthogonal to the plane of peptidoglycans. The density distributions for each peptide are shown in the insets. (C, F) Time averaged z-coordinates of amino acid residues of melittin. The color codes are according to the legends in the density profiles.

**Figure 6:**
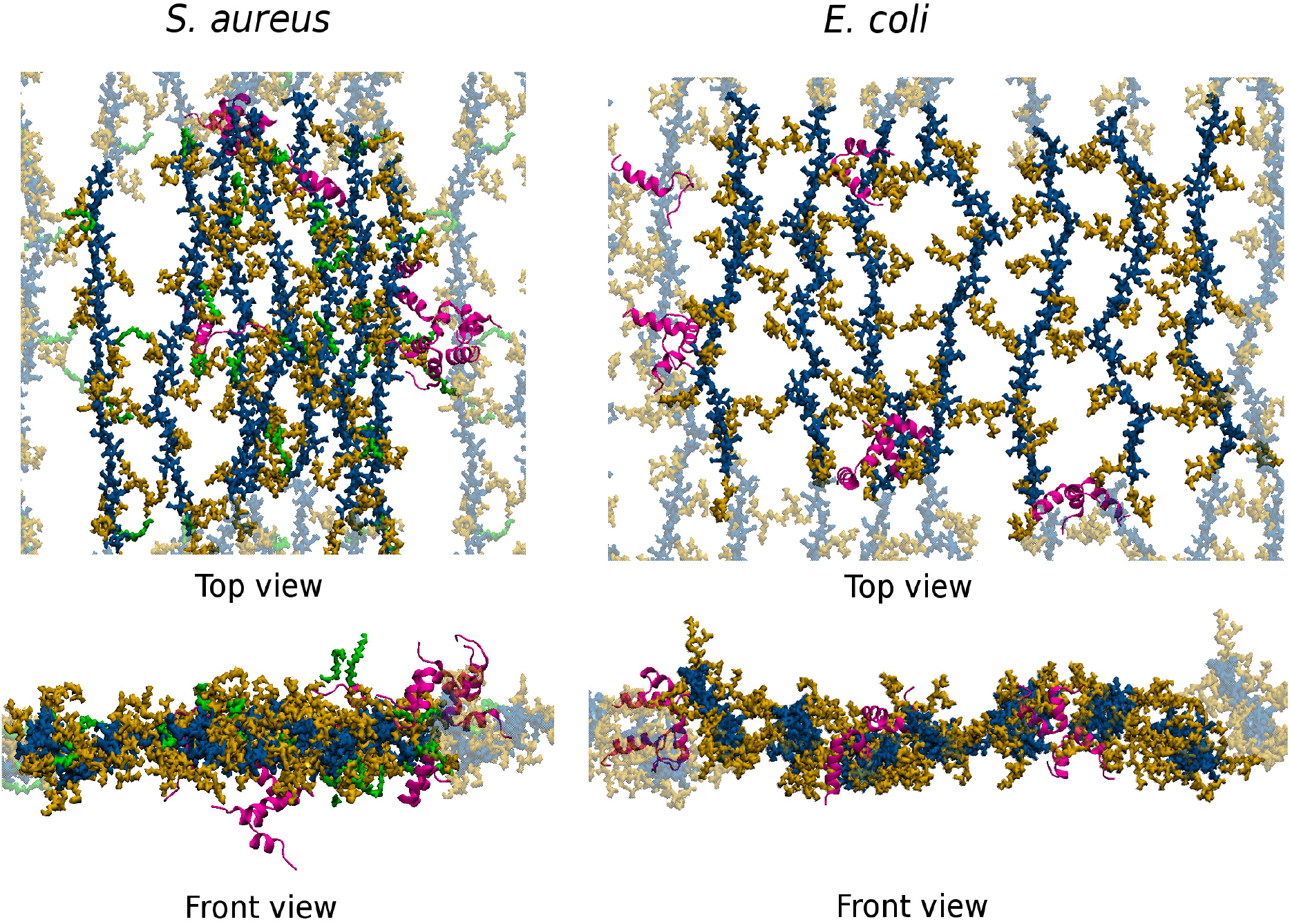
The simulation snapshots after 1*µ*s simulation of melittin peptides interacting with *S. aureus* as well as *E. coli* cell wall models. The glycan strands are shown by blue color with peptide stems in orange. The pentaglycine bridges are indicated in green, and the melittin molecules are shown in magenta. Snapshots are prepared using VMD.

We also computed the free energy landscapes (Fig. S2 in ESI) from the melittin density profiles. The energy landscapes indicate that melittin prefers to be located in the central regions of *E. coli* PGN. However, as the density profiles reveal in the case of *S. aureus*, the energy minima are present toward the surface of PGN with a small barrier of ∼1 kT present in the central regions of the membrane. These differential binding propensities for melittin between the *S. aureus* and *E. coli* membranes are also observed in the the time averaged zcoordinates of the center-of-mass of individual amino acid residues of melittins (Figs. 5C and F). Snapshots from the molecular dynamics simulations (Fig. 6) clearly illustrate the surface bound propensity of melittin for the *S. aureus* membrane. Although we did observed some tendency for aggregation of melittin on the PGN layers we did not systematically investigate this property.

The electrostatic interactions of cationic melittins with the negatively charged *E. coli* structure is expected to be stronger in comparison to the binding with the electrostatically neutral structure of the model *S. aureus*. In both the models of PGN, the melittin peptides are found to be preferentially oriented with majority of the residues interacting with PGN structures (Figs. 5C and F). The orientation was quantified by computing the helix axes according to the algorithm described elsewhere.^68^ The distributions for angles that the helix axes make with z-axis (normal to PGN) are given in Figs. S3 and S5 of ESI which indicate the preferential membrane parallel orientation adopted by melittin. We also observed that the melittin peptides retain the helicity, however some loss of helicity is observed for few peptides interacting with the PGN models (Figs. S4(G) and S6(B and D) in ESI).

From the simulations with unfolded peptides, we observed no structural transition over the 500 ns long simulations (Fig. S7 in ESI). The melittin peptides remain unfolded, indicating relatively higher stability of the unstructured state of the melittin compared to the native helix-turn-helix state.

### Melittin binds preferentially with D-Ala residues

The interactions of melittin with PGN are quantified by calculating the contacts that the melittin peptides make with PGN precursors, namely glycans (NAG and NAM) and amino acid residues of peptidoglycans. We used a distance criterion with a spherical cutoff of 0.5 nm to calculate the contacts between a melittin and a PGN segment. At a given instant, an atom of melittin making multiple contacts with a given segment of PGN is counted as unity. The contact counts are time averaged over the last 500 ns of the trajectory in each case. The counts are prominent for sugars and D-Ala residues (Fig. 7). The contacts with sugars are greater in *E. coli* as a consequence of stronger bindings with *E. coli* compared to *S. aureus*. Moreover, melittin shows a preferential binding with higher contacts for D-Ala residues among amino acids in PGN structures. This observation is in agreement with a recent experimental work on antimicrobial peptides binding to PGN, wherein melittin was shown to interact with PGN to an extent similar to vancomycin.^67^ The latter is known to bind to D-Ala moieties in PGN. ^69^ Binding of melittins with D-Ala residues is a signature of inhibitory action of melittins for transpeptidation in the biosynthesis pathway of PGN.

**Figure 7:**
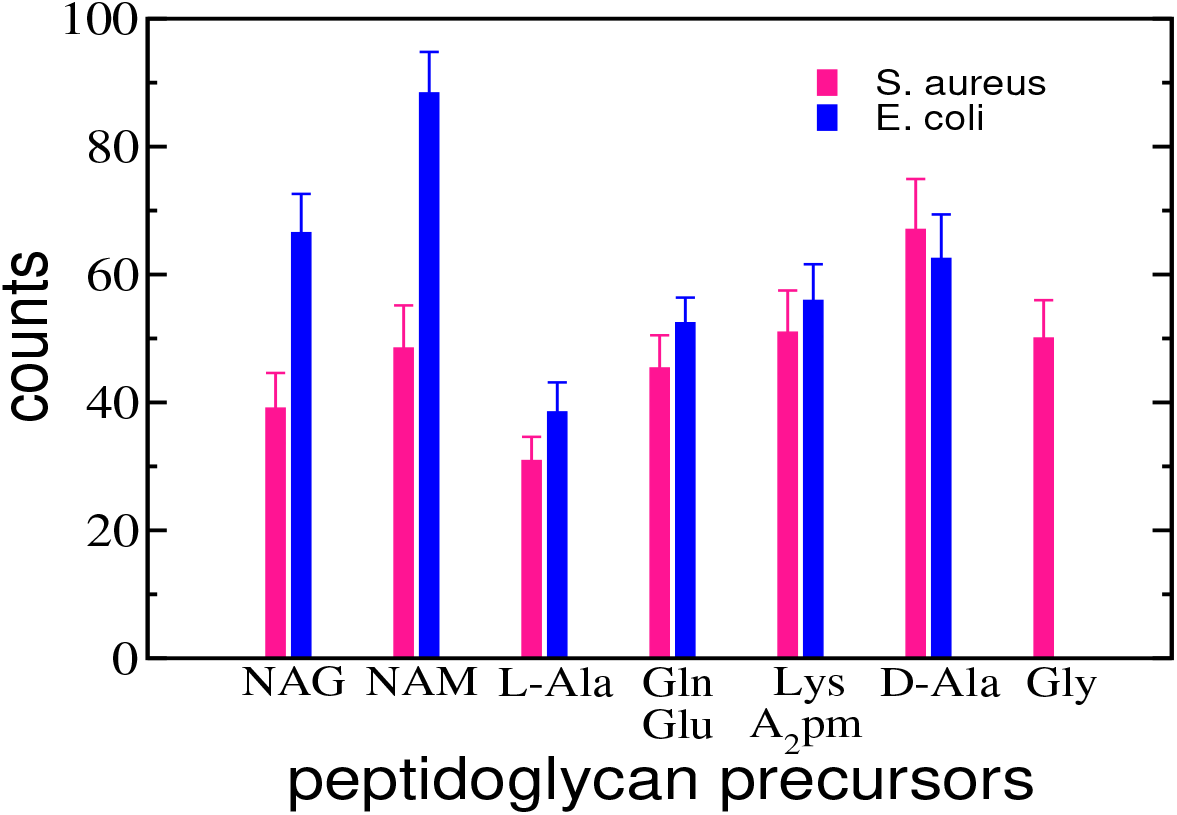
The number of contacts made by melittin with individual units of peptidoglycans for *S. aureus* and *E. coli* models. The contacts are calculated using a distance cutoff of 0.5 nm, and averaged over 500 ns long trajectories as well as averaged over the melittin peptides. The error bars represent the standard deviations.

## Barrier free energy for translocation of thymol

In addition to the interactions with melittin we also carried out molecular dynamics simulations of thymol molecules interacting with the four-layered model of *S. aureus*. Thymol is a naturally occurring antimicrobial molecule with potential use in disinfectant formations. ^70,71^ The insertion energy barriers for translocation of thymol through the complex outer and inner membranes of *E. coli* were assessed using molecular simulations in our previous studies.^72^ Figure 8(A) shows the z-component of center-of-mass of PGN layers and thymol molecules. The thymol molecules are able to translocate the multilayered PGN within the time scale of hundreds of ns, indicating the efficacy of thymol as a potential antimicrobial to translocate the thick barrier of Staphylococcal PGN. This is consistent with our previous study on *E. coli* PGN, wherein no significant barrier was observed for thymol to cross the peptidoglycan membrane.^29^ To quantify the energetics of thymol interactions, we calculated the free energy change ∆*G/kT* = − ln *ρ* using the normalized density (*ρ*) of thymol molecules. As shown in Fig. 8(B) the thymol is energetically more favourable within the PGN layers compared to aqueous environment, although the energy differences are not significant indicative of the several crossing events observed in the simulations.

**Figure 8:**
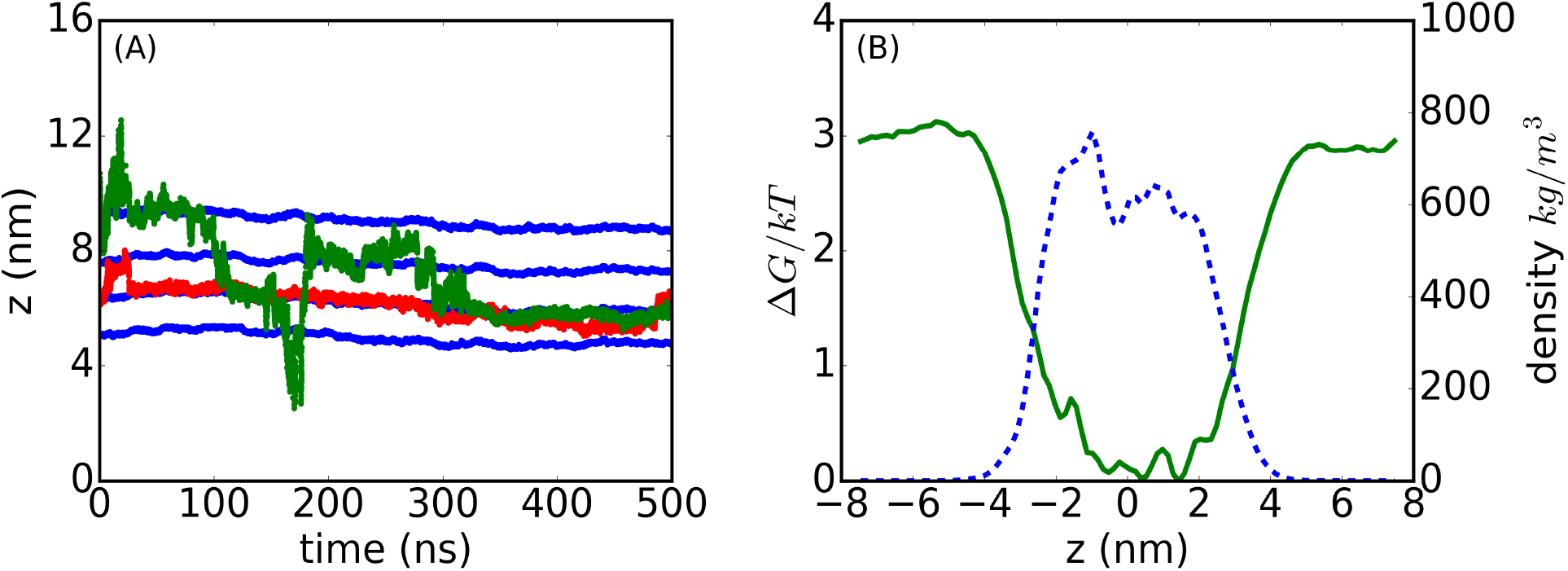
(A) The time evolution of the center-of-mass of PGN layers (blue) and two representative thymol molecules (red and green). The x-axis represents time in ns, while the y-axis shows the z-component of center-of-mass in nm. (B) Free energy of interactions of thymol with four-layered PGN structure as a function of its distance from the center of PGN (green). The density of PGN (blue) is also shown to indicate the regions of PGN.

## Conclusions

An atomistic model for the Staphylococcal cell wall peptidoglycans has been developed in this study. A detailed procedure for constructing single as well as four-layered structures of glycans and peptides for Gram-positive *S. aureus* is elucidated. The model PGN structures are able to reproduce several structural features of the cell walls of *S. aureus* bacteria. The area per disaccharide, peptide orientation, and the pore sizes are consistent with the previously reported experimental data. The Staphylococcal PGN structures are also compared with the PGN model for *E. coli*. Sharing some common features with the PGN model of *E. coli*, a linear size of ∼1 nm per disaccharide in the glycan strands is consistent with experimental data. The lateral area per disaccharide for *S. aureus* is ∼15% lower than that for the PGN of *E. coli*, indicating a relatively higher impermeability of the *S. aureus* model. A comparatively lower charge density for the PGN model of *S. aureus* is observed, and, as a consequence, the electrostatic potential is one order lower in magnitude when compared with that of PGN model for *E. coli*. We use the molecular models to unravel their interactions with melittin and classify interactions with both the PGN sugars and peptide units. Melittin is found to have a stronger interaction with the PGN of *E. coli* when compared with *S. aureus*, preferentially binding to the sugars over the peptides. Interestingly, higher atomic contacts are observed with D-Ala residues for both *S. aureus* and *E. coli* PGNs. Binding of antimicrobials with terminal D-Ala residues in cell walls is known to inhibit transpeptidation in the biosynthesis of peptidoglycans and our observations of increased binding are consistent with this finding. Molecular dynamics simulations with thymol reveals the absence of any significant barrier in the four-layered PGN model for *S. aureus*, and several translocation events were observed during the course of the simulation. Our work illustrates the utility of developing detailed atomistic models for PGN and the potential for these models to be used for in silico screening of interactions of antimicrobial molecules with the bacterial cell wall. In a departure from earlier studies a novel contribution is the development of a multilayered PGN model for Gram-positive, *S. aureus*.

## Acknowledgement

We acknowledge Thematic Unit of Excellence on Computational Materials Science (TUECMS) supported by the Department of Science and Technology (DST) for the computational facilities. We would like to thank the Supercomputer Education and Research Center (SERC) for availing computational facility at the Indian Institute of Science, Bangalore. We also thank the Department of Science and Technology (DST) and Department of Electronics and Information Technology (DeitY), India, for providing funding under the National Supercomputing Mission (NSM). We also acknowledge support from Unilever Research and Development (Bengaluru, India). We thank Morris Waskar and Janhavi Raut from Unilever for insightful discussions.

## Data Availability

The data that supports the findings of this study are available within the article and its supplementary material.

