## Supplementary material for "Differentiating interactions of antimicrobials with Gram-negative and Gram-positive bacterial cell walls using molecular dynamics simulations": SI

Electronic Supplementary Information

Rakesh Vaiwala,[a], Pradyumn Sharma,[a,c] K. Ganapathy Ayappa [a,b]

[a] *Department of Chemical Engineering, Indian Institute of Science, Bangalore 560012, India*
[b] *Centre for BioSystems Science and Engineering, Indian Institute of Science, Bangalore 560012, India*
[c] *Present address - Eli Lilly Services India Private Limited, Bangalore 560103, India*

### 1 List of molecular topology and structure files

- The single peptidoglycan chain of *S. aureus* → SA-singlechain.itp and SA-singlechain.gro
- The single layered model for *S. aureus* peptidoglycans → SA-singlelayer.itp and SA-singlelayer.gro
- The four-layered model for *S. aureus* peptidoglycans → SA-fourlayer.itp and SA-fourlayer.gro
- The single peptidoglycan chain of *E. coli* → EC-singlechain.itp and EC-singlechain.gro
- The single layered model for *E. coli* peptidoglycans → EC-singlelayer.itp and EC-singlelayer.gro

### 2 List of Figures

S1. Native structure of melittin and the its secondary structure in water.

S2. Free energy profiles for melittin interacting with the model *S. aureus* and *E. coli* cell wall structures.

S3. The probability histograms for the angles by helix axes with z-axis for melittins interacting with the model *S. aureus* cell wall.

S4. The secondary structure analysis for melittin during the course of interactions with the PGN structure of *S. aureus*.

S5. The probability histograms for the angles made by helix axes with z-axis for melittins interacting with the model *E. coli* cell wall.

S6. The secondary structure analysis for melittin during the course of interactions with the PGN structure of *E. coli*.

S7. The secondary structure of unfolded melittin peptides during their interactions with single layered PGN models of *S. aureus* and *E. coli*.

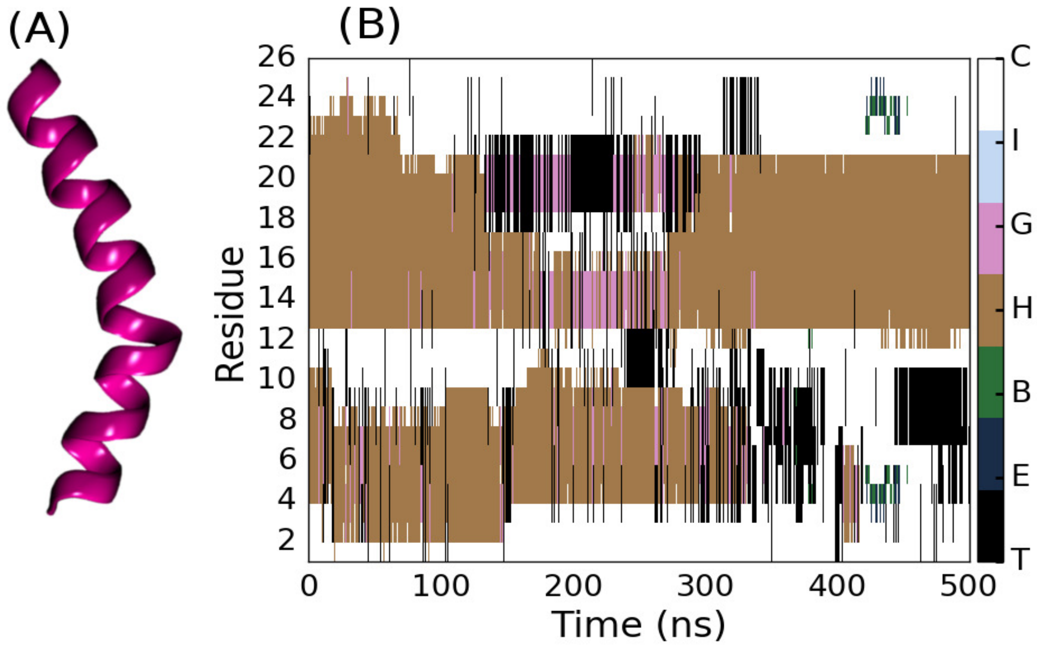


**Figure S1:** (A) Native structure of melittin (PDB: 2MLT). (B) Secondary structure analysis for melittin peptide simulated in water. Melittin tends to lose helicity in aqueous environment. The color bar represents structures - turns T, extended conformations E, isolated bridge B, $\alpha$-helix H, 3-10 helix G, $\pi$-helix I and coil C.

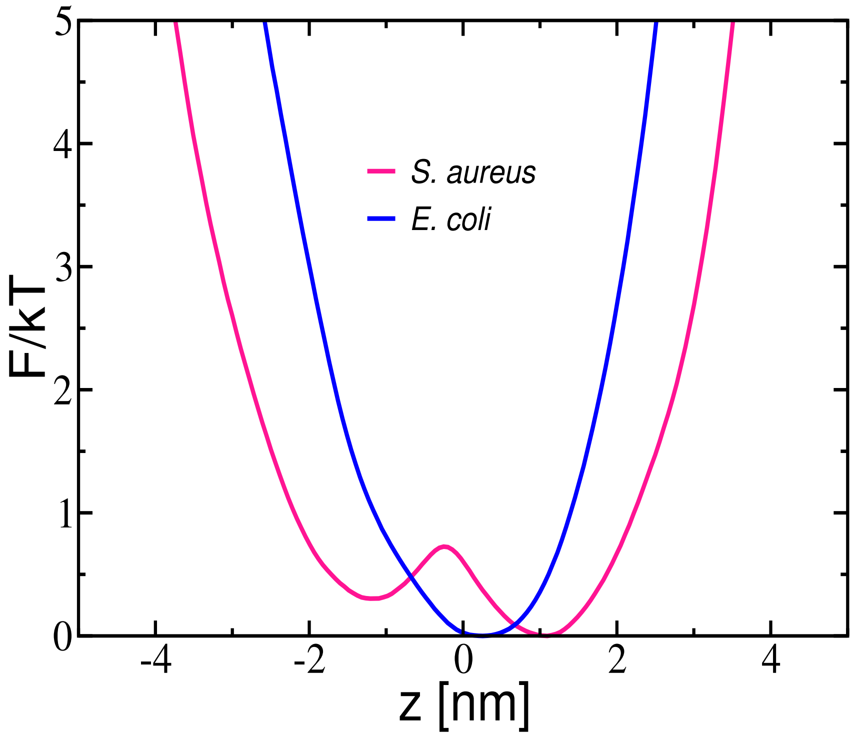


**Figure S2:** Free energy profiles for melittin interacting with the model *S. aureus* and *E. coli* cell wall structures. The energies F/kT = $-\log(\rho/\rho_{max})$ are derived from density distributions for melittin given in Figures 5B and 5E of the main report.

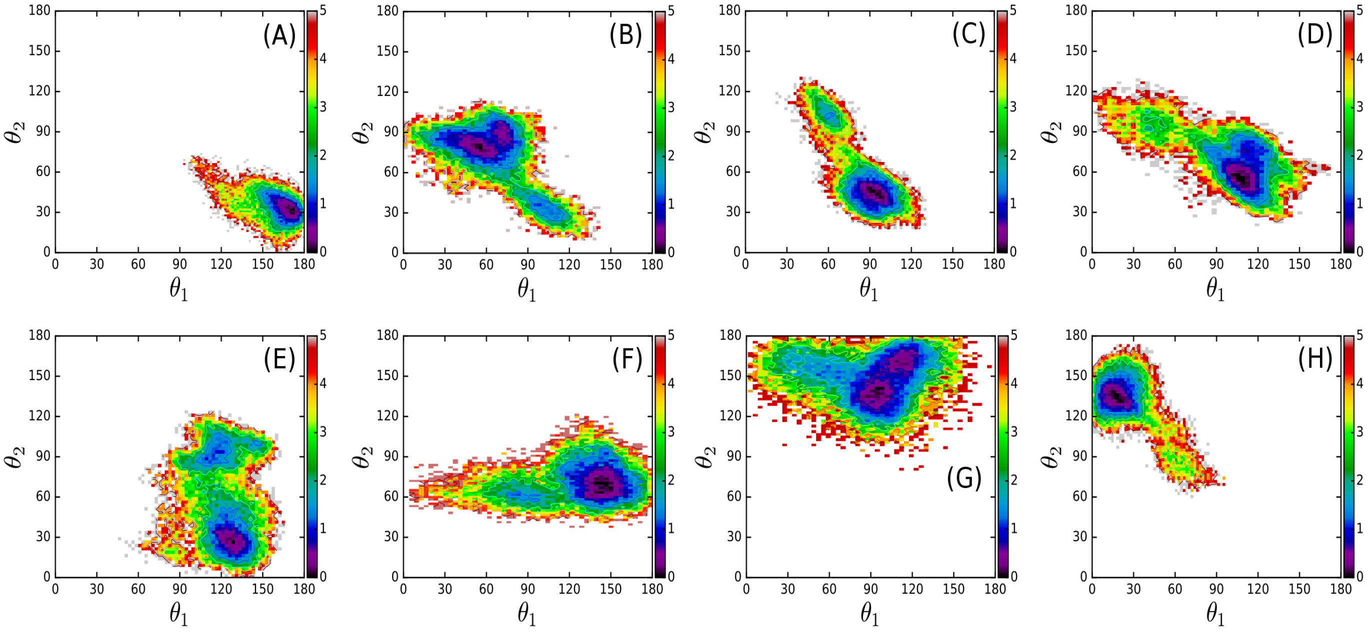


**Figure S3:** The joint probability histograms $H$ for the angles made with z-axis by the helix axes in melittin interacting with the model *S. aureus* cell wall. The helix axes computed (Ref. Kahn Peter C., *Computers & Chemistry* (1989) 13(3) 185-189) using residues GLY1-THR11 at N-terminus and PRO14-GLN26 at C-terminus, defining the angles $\theta_1$ and $\theta_2$, respectively. The coordinates of $C_\alpha$ atoms are used to compute the helix axes. The color map is the log-normal scale $-\log(H/H_{max})$, with $H_{max}$ being the maximum value of $H$ in histograms. The sequence A-H corresponds to the peptides 1-8 given in Figure 5B of the main report.

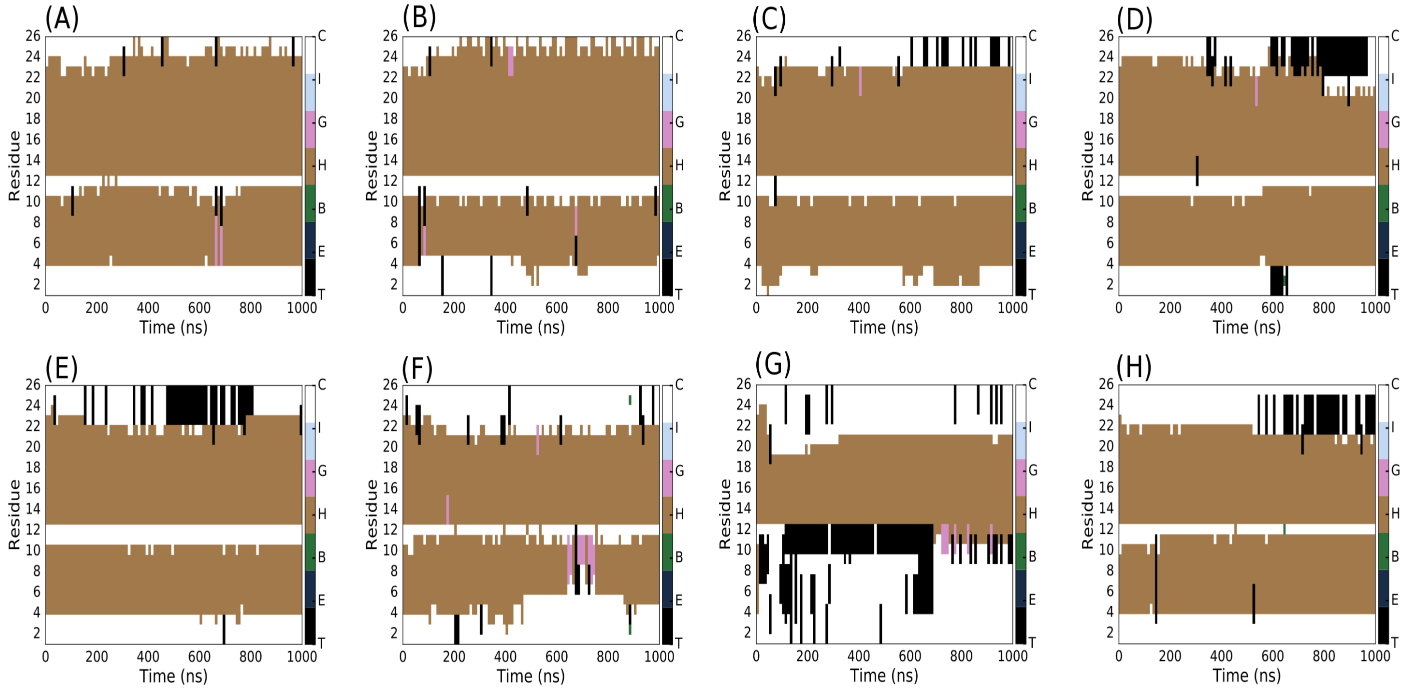


**Figure S4:** The secondary structure analysis for melittin during the course of interactions with the PGN structure of *S. aureus*. The panel sequence (A-H) corresponds to the melittin peptides 1-8 given in Figure 5B of the main report. The color bar represents structures - turns T, extended conformations E, isolated bridge B, $\alpha$-helix H, 3-10 helix G, $\pi$-helix I and coil C.

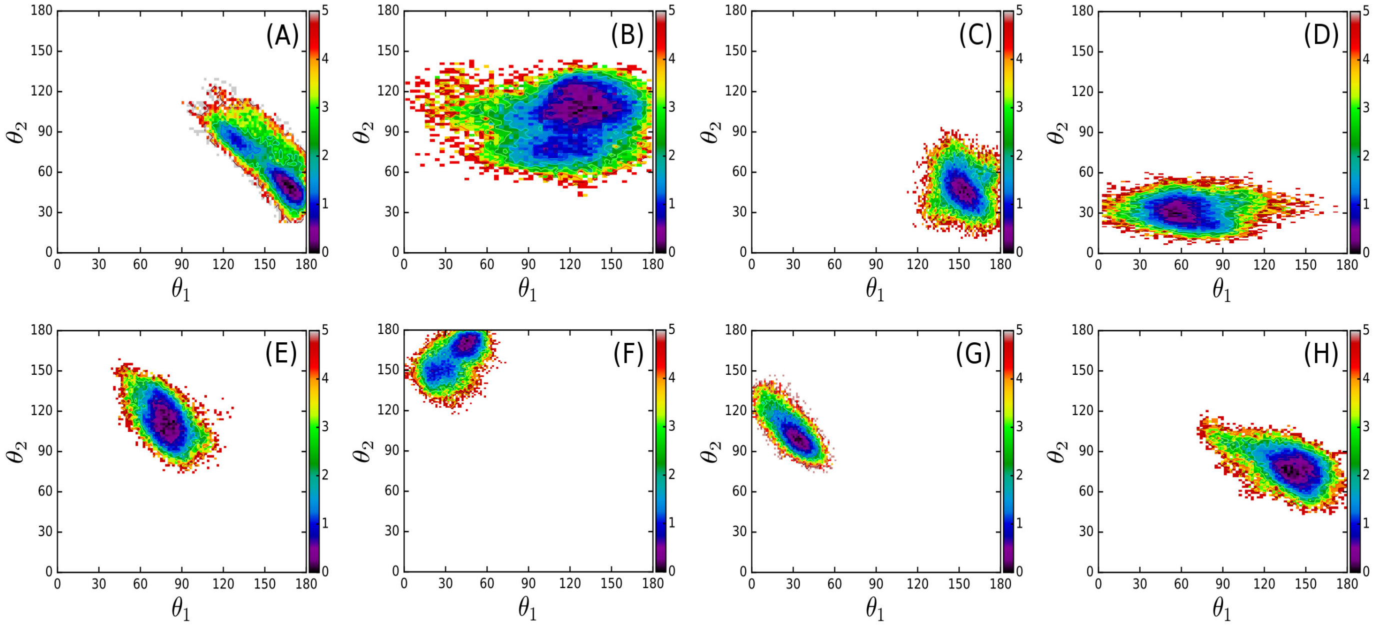


**Figure S5:** The joint probability histograms $H$ for the angles made with z-axis by the helix axes in melittin interacting with the model *E. coli* cell wall. The helix axes computed using residues GLY1-THR11 at N-terminus and PRO14-GLN26 at C-terminus make angles $\theta_1$ and $\theta_2$, respectively. The coordinates of $C_\alpha$ atoms are used to compute the helix axes. The color map is the log-normal scale $-\log(H/H_{max})$, with $H_{max}$ being the maximum value of $H$ in histograms. The sequence A-H corresponds to the melittin peptides 1-8 given in Figure 5E of the main report.

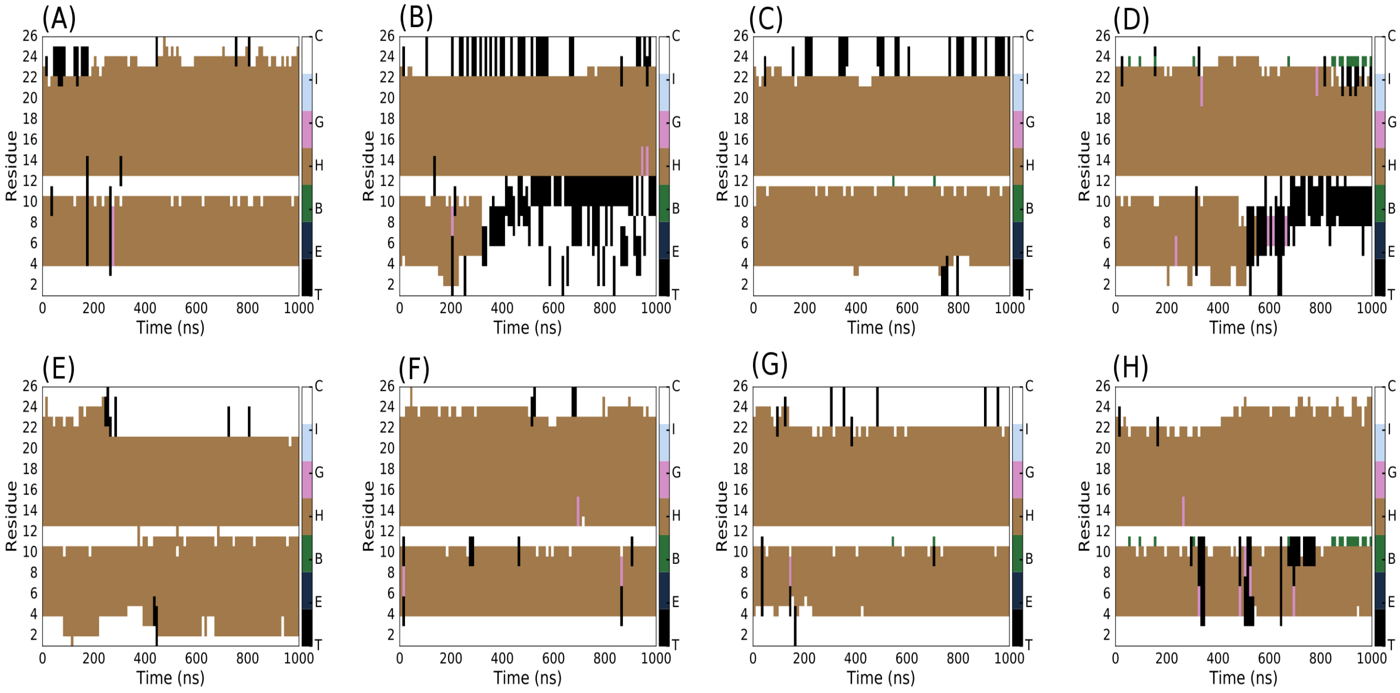


**Figure S6:** The secondary structure analysis for melittin during the course of interactions with the PGN structure of *E. coli*. The panel sequence (A-H) corresponds to the melittin peptides 1-8 given in Figure 5E of the main report. The color bar represents structures - turns T, extended conformations E, isolated bridge B, $\alpha$-helix H, 3-10 helix G, $\pi$-helix I and coil C.

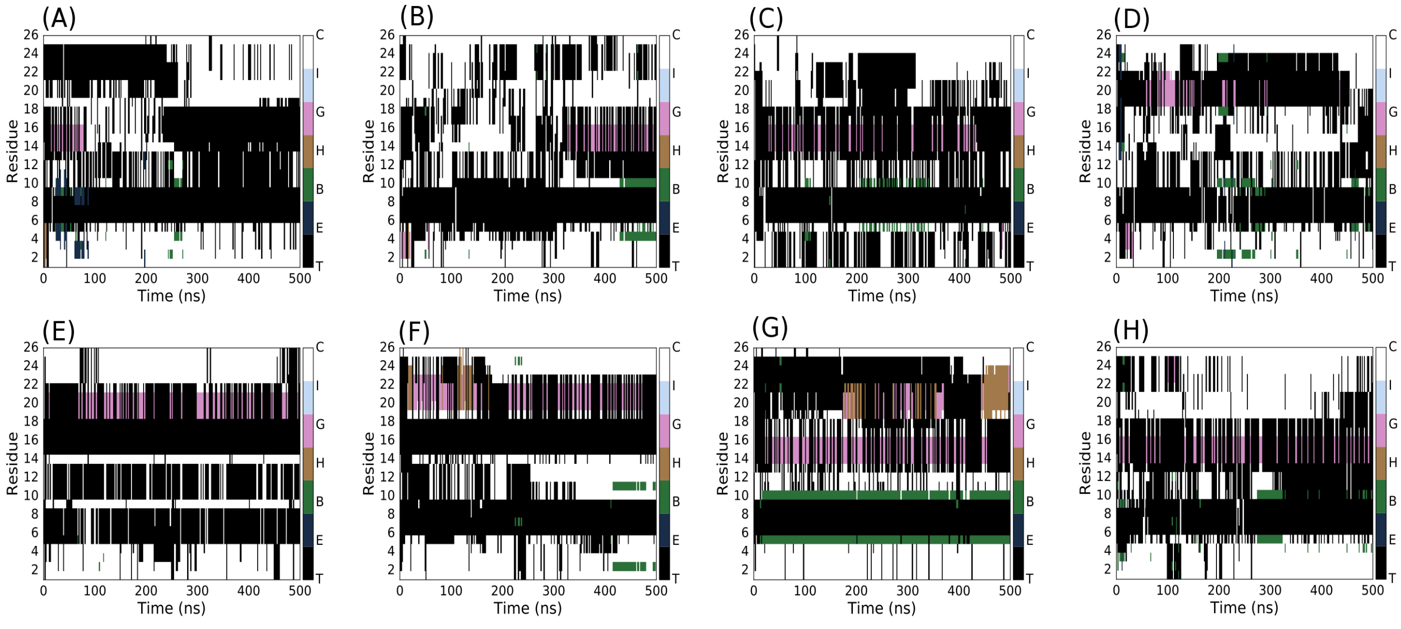


**Figure S7:** The time evolution of secondary structure of unfolded melittin peptides during interactions with single layered models of *S. aureus* (A-D) and *E. coli* (E-H) cell walls.
